# Recombinant production of active microbial transglutaminase in *E. coli* by using self-cleavable zymogen with mutated propeptide

**DOI:** 10.1101/2020.02.25.965541

**Authors:** Ryo Sato, Kosuke Minamihata, Ryutaro Ariyoshi, Hiromasa Taniguchi, Noriho Kamiya

**Affiliations:** Department of Applied Chemistry, Graduate School of Engineering, Kyushu University, 744 Motooka, Fukuoka 819-0395, Japan; Division of Biotechnology, Center for Future Chemistry, Kyushu University, 744 Motooka, Fukuoka 819-0395, Japan

**Keywords:** microbial transglutaminase, *Streptomyces mobaraensis*, zymogen, self-cleavable, chimera protein

## Abstract

Microbial transglutaminase from *Streptomyces mobaraensis* (MTG) has been widely used in food industry and also in research and medical applications, since it can site-specifically modify proteins by the cross-linking reaction of glutamine residue and the primary amino group. The recombinant expression system of MTG in *E. coli* provides better accessibility for the researchers and thus can promote further utilization of MTG. Herein, we report production of active and soluble MTG in *E. coli* by using a chimeric protein of tobacco etch virus (TEV) protease and MTG zymogen. A chimera of TEV protease and MTG zymogen with native propeptide resulted in active MTG contaminated with cleaved propeptide due to the strong interaction between the propeptide and catalytic domain of MTG. Introduction of mutations of K9R and Y11A to the propeptide facilitated dissociation of the cleaved propeptide from the catalytic domain of MTG and active MTG without any contamination of the propeptide was obtained. The specific activity of the active MTG was 22.7±2.6 U/mg. The successful expression and purification of active MTG by using the chimera protein of TEV protease and MTG zymogen with mutations in the propeptide can advance the use of MTG and the researches using MTG mediated cross-linking reactions.

## Introduction

Transglutaminase (TGases, protein-glutamine γ-glutamyltransferase, E.C. 2.3.2.13) catalyzes acyl transfer reaction between γ-carboxyamide groups of glutamine residues and primary amine including ε-amino groups of lysine residues. Among many transglutaminases from different sources [1], a microbial transglutaminase from *Streptomyces mobaraensis* (MTG) is the one studied the most and is commercially available today [2]. Although the main application of MTG is in food industry [3,4], MTG has been used as a ligation tool for proteins in research [5] and pharmaceutical fields [6]. MTG-mediated site-specific conjugations of proteins with small functional molecules [7], protein [8], DNA [9,10], RNA[11], synthetic polymers [12], and lipids [13] have been reported. Moreover, MTG is utilized to conjugate anticancer agents to antibody to make antibody-drug conjugates (ADCs) for medical use [14,15]. For those researches and uses, the precise control of reaction condition is necessary to achieve site-specific modifications of proteins and correct assessment of the obtained results and thus a highly purified MTG is needed. The commercial MTG products for food industry are cheap, however, they contain a lot of additives as well as other proteinous substances, and thus further purification is required before use. A commercial MTG with high purity and activity is available from Zedira (Germany), yet its price is rather expensive for routine use. Therefore, it is meaningful to develop an *E. coli* expression system of MTG for laboratory use with high purity and activity.

In nature, MTG is produced as a zymogen, carrying a propeptide at the N-terminus of matured MTG domain and proteolysis of the propeptide is required for activation of MTG [16]. The propeptide has a function of intramolecular chaperone and thus the expression of MTG without the propeptide in *E. coli* results in the formation of inclusion body [17]. The active MTG can be made by refolding from the inclusion body [18] or by removing the propeptide from MTG zymogen by protease treatment [19]. However, these strategies are not straightforward, because additional refolding process or protease treatment process as well as another purification steps after proteolysis are needed.

One step expression of the active MTG in *E. coli* has been demonstrated by polycistronic expression of the propeptide and the matured MTG domain in *E. coli* [20,21]. Both were secreted into the periplasmic space and the propeptide facilitated the folding of matured MTG domain and active MTG was obtained in soluble form without any additional downstream processes. Another approach was to co-express MTG zymogens with a protease [22]. MTG zymogens with 3C protease recognition sequence between the propeptide and matured MTG domain were successfully processed by the co-expressed 3C protease in *E. coli* and active MTG was obtained. For both strategies, the expression level of two genes would have impact on the overall yield of active MTG. The simplest but the most precise way to control the expression ratio of two proteins is to genetically fuse and express them as a single polypeptide.

Herein we report another strategy to make active MTG in soluble form by constructing a chimera protein of a protease and MTG zymogen. Tobacco etch virus (TEV) protease was selected to cleave off the propeptide from MTG zymogen. Two constructs of chimera proteins of TEV protease and MTG zymogen were constructed by using pMAL-c5E vector (**Fig. 1A**). Both constructs contain maltose-binding protein (MBP) at the N-termini to promote the correct folding of the chimera proteins and to increase the solubility of them. The N-terminal amino group of Gly can be a substrate of MTG reaction [23]. Therefore, to prevent self-cross-linking of MTG after propeptide cleavage by TEV protease, a TEV protease recognition sequence of GSENLYFQ↓SGG was inserted between the propeptide and matured MTG domain in MBP-TEV-Pro-MTG (**Fig. 1B**). It’s been reported that the modulation of interaction between propeptide and the catalytic domain of MTG was key to increase the productivity of active MTG [22]. Moreover, in this study, we put another mutation of K9R in the propeptide to eliminate possibility of cross-linking of MTG zymogen via K9. MBP-TEV-Pro(K9R/Y11A)-MTG has mutations of K9R and Y11A in the propeptide and a longer TEV protease recognition sequence (GGGSENLYFQSGGGGS) than MBP-TEV-Pro-MTG with deleting 8 amino acid residues (AGPSFRAP) at the C-terminal of propeptide (**Fig. 1C**) The whole amino acid sequences of constructed MTG variants were provided in Supplementary Information as **Table S1**. We anticipated the mutations in the propeptide attenuate the interaction towards the catalytic domain of MTG and facilitate dissociation of the propeptide from active-MTG after self-cleavage (**Fig. 1D**).

**Fig. 1.**
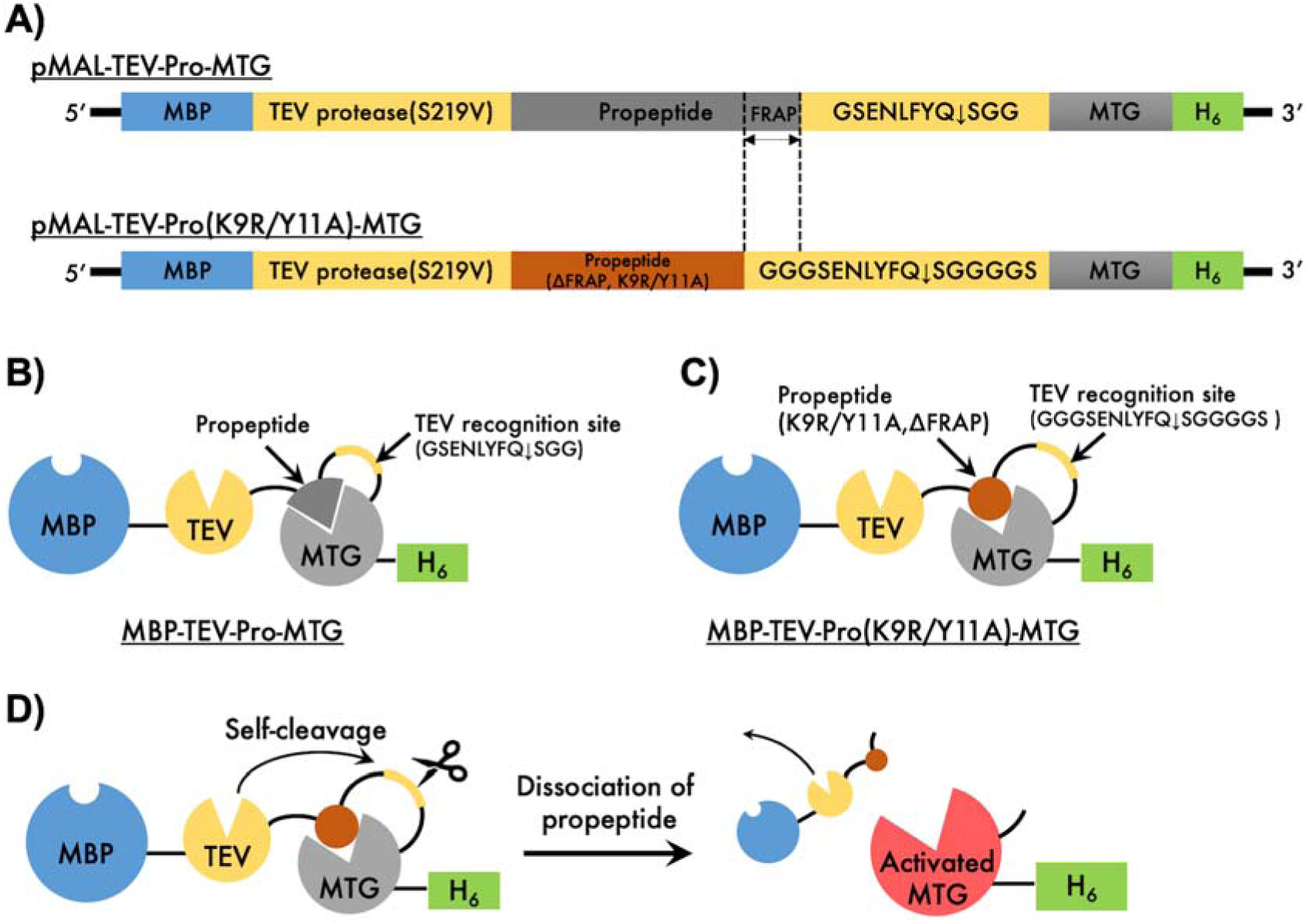
Constructs of MTG variants evaluated in this study. A: Schematic illustrations of DNA constructs of pMAL-TEV-Pro-MTG and pMAL-TEV-Pro(K9R/Y11A)-MTG. B and C: Schematic illustrations of MBP-TEV-Pro-MTG and MBP-TEV-Pro(K9R/Y11A)-MTG, respectively. D: Self-cleavage reaction of MBP-TEV-Pro(K9R/Y11A)-MTG to obtain fully activated MTG.

## Material and Methods

### 2.1 Materials

PrimeSTAR® MAX DNA Polymerase and In-Fusion® HD Cloning Kit were purchased from Takara Bio Inc (Shiga, Japan). LB broth (Miller) was purchased from Merck Millipore Corp (Billerica, MA, USA). Tryptone, extract yeast dried, tris (hydroxymethyl) aminomethane (Tris), acrylamide/bis mixed solution (29:1), and Rapid Stain CBB kit were purchased from Nacalai Tesque, Inc. (Kyoto, Japan). Prestained XL-Ladder was purchased from APRO Science Inc (Tokushima, Japan). Hydrochloric acid, sodium chloride, potassium chloride, disodium hydrogenphosphate 12-water, dipotassium hydrogenphosphate, potassium dihydrogen phosphate, glutathione (reduced form), Fe(III) chloride hexahydrate, and trichloroacetic acid were purchased from FUJIFILM Wako Pure Chemical Corporation (Osaka, Japan). N,N,N’,N’-tetramethylethylenediamine, ammonium persulfate, and hydroxylamine hydrochloride were purchased from Kishida Chemical Co.,Ltd (Osaka, Japan). Z-QG was purchased from Peptide Institute, Inc. (Osaka, Japan). L-glutamic acid γ-monohydroxamate was purchased from Sigma-Aldrich Co. LLC (St. Louis, MO). Ultrapure water supplied by Milli-Q® Reference Water Purification System (Merck Millipore Corp., Billerica, MA, USA) was used for buffer preparation.

### 2.2 Construction of expression vectors and preparation of recombinant proteins

The gene encoding tobacco etch virus (TEV) protease mutant S219V [24] was inserted at the 3’ end of gene encoding maltose binding protein (MBP) in pMAL-c5E vector (New England Biolabs, MA, USA) to construct pMAL-TEV vector. A synthetic gene encoding pro-MTG from *Streptomyces mobaraensis*, which has C-terminal HisTag sequence (H_6_) and TEV protease recognition sequence and linkers (GSENLYFQSGG) at between the propeptide and catalytic domain of MTG was then fused to the 3’ end of the gene of TEV protease in the pMAL-TEV vector to construct pMAL-TEV-Pro-MTG. Two mutations at K9R and Y11A in the propeptide of MTG were then introduced to pMAL-TEV-Pro-MTG. Also the linker sequence at TEV protease recognition site was extended (GGGSENLYFQSGGGGS) with deleting 8 amino acid residues (AGPSFRAP) at the C-terminal of propeptide to construct pMAL-TEV-Pro(K9R/Y11A)-MTG. (**Figure 1**) The whole amino acid sequences of constructed MTG variants were provided in **Table S1**.

The expression of MTG variants was conducted on an *Escherichia coli* BL21 Star (DE3) strain. One ng of the expression vectors of MTG variants were transformed into the cells by a heat shock method and cultured on a LB agar medium containing 100 μg/mL of ampicillin sodium. A single colony was inoculated to 5.0 mL of LB medium containing the same amount of ampicillin sodium and shaken at 220 rpm at 37 °C for 6 hours. One ml of the precultured medium was added into 250 mL of Terrific broth (TB) medium containing the same antibiotics and cultured with shaking at 220 rpm at 37 °C until the optical density at 600 nm reached around 0.5. TB medium contains 3 g of tryptone, 6 g of yeast extract, 2 mL of glycerol, 2.35 g of dipotassium hydrogen phosphate and 0.55 g of potassium dihydrogen phosphate in 250 mL. Expression of each protein was induced by addition of isopropyl β-d-1-thiogalactopyranoside at a final concentration of 0.1 mM, and the cells were cultured at 17°C for 16 hours. The cells were harvested by centrifugation at 5,000 g for 20 min and supernatant was removed. The cell pellet was resuspended in PBS (-) (pH 7.4) and wet weight of each pellet was measured after removing PBS(-). The expression in TB medium was conducted in triplicate for each MTG variants.

For the purification of MTG variants, the cells were resuspended again in PBS(-) and the cells were disrupted by sonication on ice for 12.5 min. The cell debris was removed by centrifugation at 18,000g for 20 min at 4 °C. The supernatant was initially applied to a HisTrap FF crude 5 mL (GE Healthcare UK Ltd.), which were pre-equilibrated with the HisTrap binding buffer (20 mM Tris-HCl, 500 mM NaCl, 35 mM imidazole, pH 7.4). The column was washed with the same buffer until all the unbound proteinaceous substances eluted out. The MTG variants were eluted by a gradient of HisTrap elution buffer (20 mM Tris-HCl, 500 mM NaCl, 500 mM imidazole, pH 7.4) up to 100% with 13 column volumes. The fractions containing MTG variants were concentrated to a volume of approximately 5 mL using ultrafiltration membrane of 10 kDa MWCO (Vivaspin Turbo 15, Sartorius). Subsequently, MTG variants were purified with size exclusion chromatography using HiLoad 16/600 Superdex 75 pg (GE Healthcare UK Ltd.) with PBS (pH 7.4) containing 0.1 mM of dithiothreitol as a running buffer. The MTG variants were concentrated using the 10 kDa MWCO ultrafiltration membrane. The protein concentrations of purified MTG variants were determined by measuring absorbance at 280 nm using an ND-1000 Nanodrop and calculated by using the extinction coefficient of MTGs predicted by ProtParam at ExPASy websites.

### 2.3 Activity measurement of the active MTG by hydroxamate assay

The activities of MTG variants were assessed by quantifying the amount of hydroxamate resulted from MTG reaction using complex formation of hydroxamate and Fe(III) ions [25]. The substrate solution containing 30mM Z-QG, 100 mM hydroxylamine hydrochloride, 10 mM glutathione in 0.2 M Tris-Acetate (pH 6.0) was added to MTG variants. After 10 min of incubation at 37 °C, the stop solution (200 mM trichloroacetic acid in 50 mM HCl) containing 50 mM of Fe(III) chloride hexahydrate was added to the reaction mixture and the absorbance at 525 nm was measured using Synergy HTX Multi-Mode Reader (BioTek, Winooski, USA). A standard curve was made by using L-glutamic acid γ-monohydroxamate solution with known concentration. One unit of MTG was defined as the amount of MTG catalyzes formation of 1 µmol of hydroxamic acid in one minute.

## 3. Results and Discussions

### 3.1 Active and soluble MTG purification

Both MTG constructs were expressed in *E. coli* BL21(DE3) in 250 mL of TB medium and the wet weights of cell pellets obtained after expression were measured in triplicate. The average wet weights of cell pellets of MBP-TEV-Pro-MTG and MBP-TEV-Pro(K9R/Y11A)-MTG were 4.2 ±0.1 g and 1.0 ±0.1 g, respectively. Introduction of mutations to the propeptide significantly reduced the amount of *E. coli* cells, suggesting that the MTG exhibited its catalytic activity in the cells and resulted in toxic effect. The MTG variants were purified with HisTag affinity chromatography and size-exclusion chromatography (SEC) and the eluates were analyzed with SDS-PAGE (**Fig. 2**). In case of MBP-TEV-Pro-MTG, the bands of the active MTG and cleaved MBP-TEV-Pro were appeared around 38 kDa and 70 kDa, respectively (**Fig. 2A**). The cleaved MBP-TEV-Pro, which lacks HisTag, bound to Ni-NTA column by attaching to the active MTG because of the strong affinity between them. On the other hand, MBP-TEV-Pro(K9R/Y11A)-MTG showed mainly the band of active MTG at around 38 kDa (**Fig. 2B**) and the most of cleaved MBP-TEV-Pro(K9R/Y11A) could be removed from the active MTG. The remaining portion of MBP-TEV-Pro(K9R/Y11A) and highly cross-linked proteins were successfully removed from the active MTG by following SEC purification (**Fig. 2E and 2F**). In contrast, in the SEC purification of MBP-TEV-Pro-MTG, all the active MTG was eluted with the cleaved MBP-TEV-Pro and highly cross-linked proteins (**Fig. 2C and 2D**) and it was impossible to isolate the active MTG. From these results, we concluded that in order to obtain active MTG without any contamination of the propeptide domain, introduction of mutations to the propeptide is essential. Some portions of active MTG were eluted with the cross-linked proteins in SEC purification of MBP-TEV-Pro(K9R/Y11A)-MTG and this is probably due to the avidity effect of the propeptide in the highly cross-linked proteins, which is composed of probably MBP-TEV-Pro(K9R/Y11A) and other endogenous proteins in *E. coli*. We expect that the productivity of MTG can be improved more by reducing the formation of such highly cross-linked proteins by eliminating the cross-linking sites of MBP-TEV-Pro(K9R/Y11A). Note that the purified active MTG in this study showed no self-cross-linking behavior (**Fig. S1**), which is reported in the previous literature [22].

**Fig. 2.**
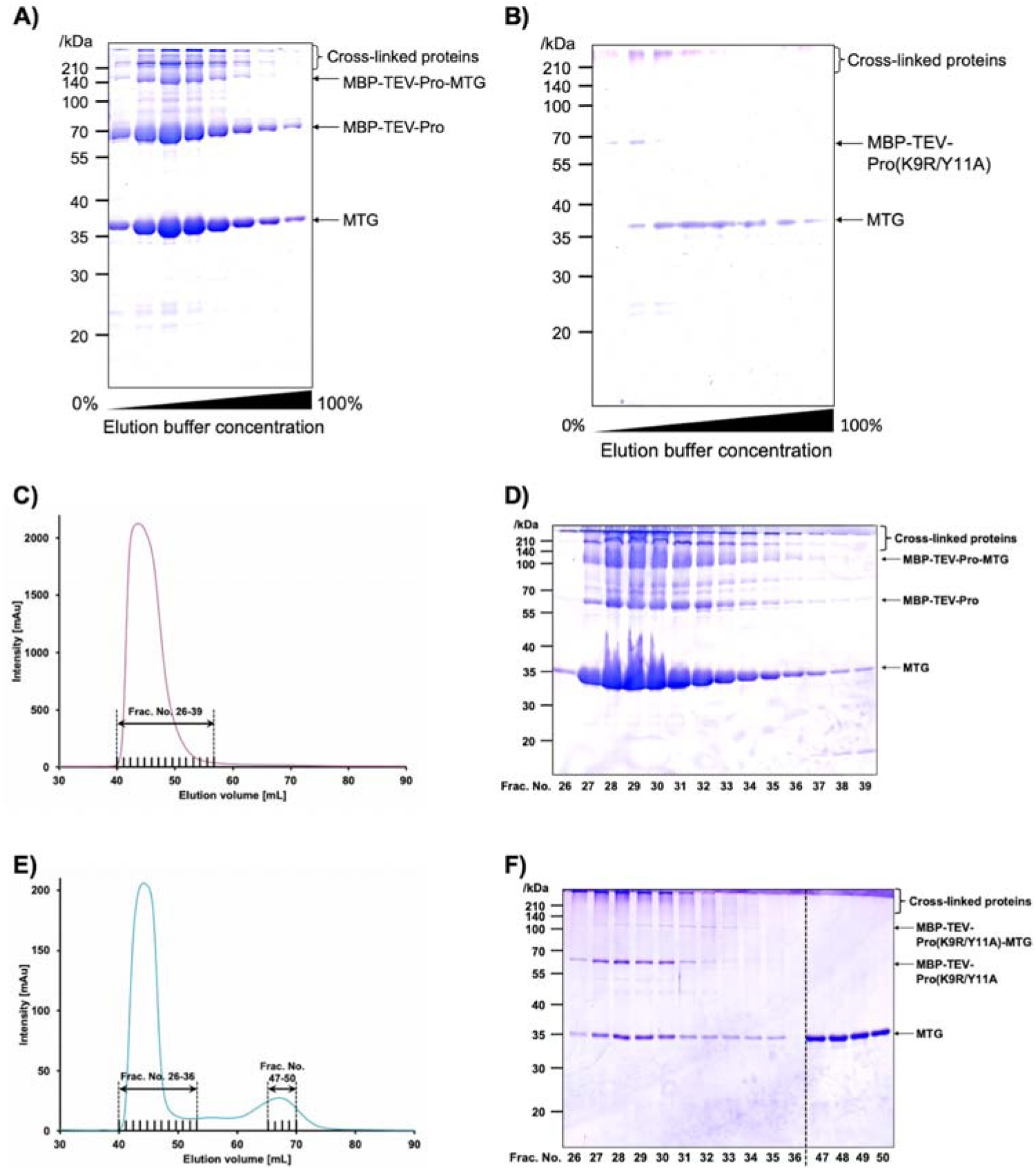
Purification of active MTG by using constructs of MBP-TEV-Pro-MTG and MBP-TEV-Pro(K9R/Y11A)-MTG. A and B: SDS-PAGE analyses of fractions in the HisTag purification of MBP-TEV-Pro-MTG and MBP-TEV-Pro(K9R/Y11A)-MTG, respectively. The elution buffer used was 25 mM Tris-HCl, pH7.4, 500 mM NaCl, 500 mM imidazole. C and D: SEC chromatogram and the SDS-PAGE analysis of fractions indicated in the chromatogram of MBP-TEV-Pro-MTG after HisTag purification (A). E and F: SEC chromatogram and the SDS-PAGE analysis of fractions indicated in the chromatogram of MBP-TEV-Pro(K9R/Y11A)-MTG after HisTag purification (B).

### 3.2 Activity measurement of MTG mutant

After collection of fractions containing MTGs (Frac. No. 26∼39 for MBP-TEV-Pro-MTG and Frac. No. 57∼50 for MBP-TEV-Pro(K9R/Y11A)-MTG shown in Fig. 2), MTG activity measurement was conducted by hydroxamate assay [25]. One unit of MTG activity was defined as the amount of enzyme that catalyzes the formation of 1 µmol of hydroxamate in 1 minute. The specific activities of the active MTGs prepared from MBP-TEV-Pro-MTG and MBP-TEV-Pro (K9R/Y11A)-MTG were 4.0±0.1 U/mg and 22.7±2.6 U/mg, respectively (**Fig. 3**). The active MTG obtained from MBP-TEV-Pro (K9R/Y11A)-MTG showed 5-fold higher specific activity than the one from MBP-TEV-Pro-MTG, because the cleaved propeptide, which has inhibitory effect, remained in the sample prepared from MBP-TEV-Pro-MTG. The specific activity of MTG from MBP-TEV-Pro (K9R/Y11A)-MTG in this study was as high as the activity of previously reported active MTG purified from the original strain [26], prepared by in vitro activation of MTG zymogen (Chen et al., 2013) and also that of the commercially available MTG (Zedira, Germany) (25 U/mg). The total amount of active MTG obtained from MBP-TEV-Pro(K9R/Y11A)-MTG was 0.23 mg from 750 mL of TB medium. Since the productivity of MTG in *E. coli* and its activity are in the relationship of trade-off, the amount of active MTG obtained in this study was not high enough for industrial production. However, it is sufficient for use in laboratories and moreover the quality and the specific activity of the obtained MTG were suitable for research uses, which require precise control of reactions.

**Fig. 3.**
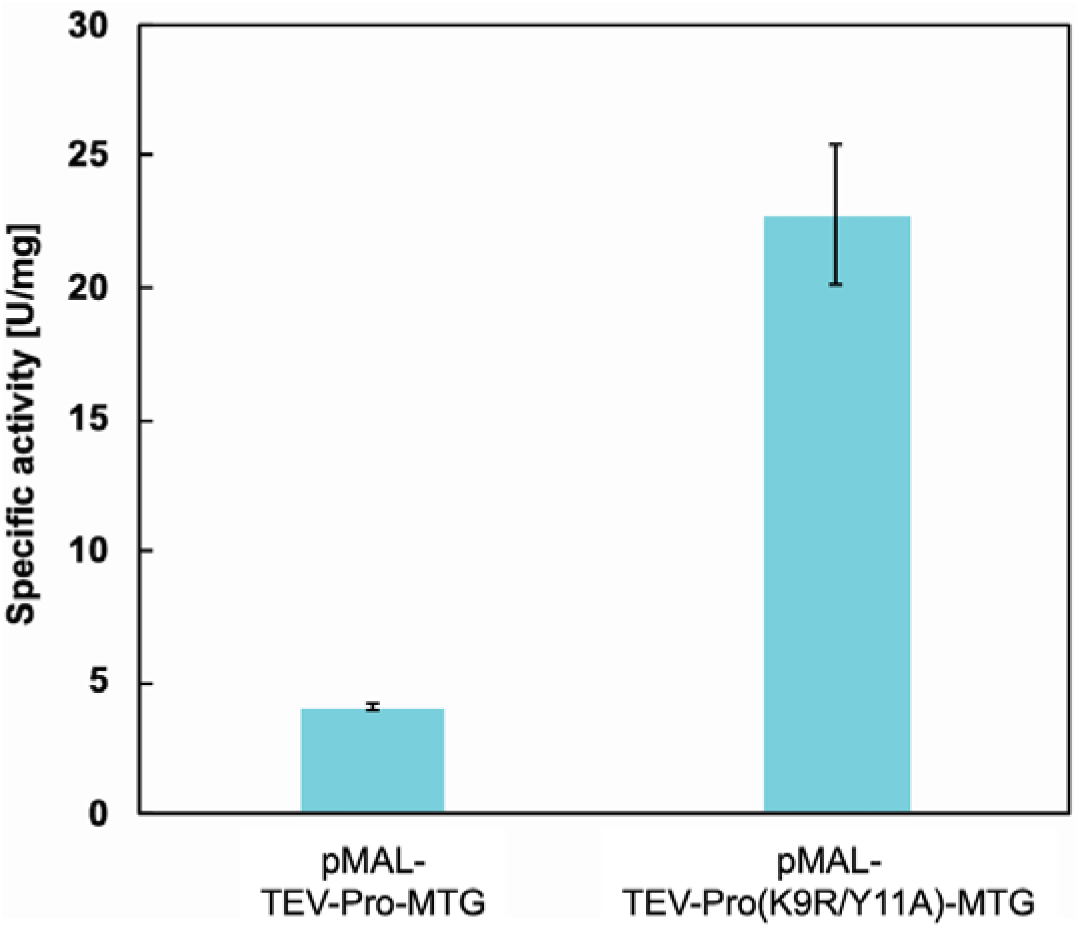
Specific activities of the active MTG prepared from two constructs of MTGs measured by hydroxamate assay.

## Conclusion

In conclusion, we demonstrated production of active and soluble MTG by constructing a chimeric protein of TEV protease and MTG zymogen. Introduction of mutations of K9R and Y11A to the propeptide was required to obtain active MTG free of contamination of cleaved propeptide, resulting in successful production of active MTG with a high specific activity.

## Acknowledgments

This work was supported by the Japan Society for the Promotion of Science (JSPS) KAKENHI Grant number JP19H00841 (to N. K.).

## Conflict of interest

The authors declare no financial or commercial conflict of interest.

## Appendix A. Supplementary data

Supplementary material related to this article can be found, in the online version, at doi:

## Abbreviations^1^

MTG: Microbial Transglutaminase;
TEV: Tobacco Etch Virus; Maltose-Binding Protein;
SEC: Size-Exclusion Chromatography

## Notes

### Competing Interest Statement

The authors have declared no competing interest.

### Summary of Updates

After being rejected by a journal, we revised our manuscript and reformatted for submission to other journal.

