## Supplementary information for "Recombinant production of active microbial transglutaminase in *E. coli* by using self-cleavable zymogen with mutated propeptide"

**Table S1.** Amino acid sequence of MTG constructs prepared in this study

| **Proteins** | **Sequences** |
| --- | --- |
| MBP-TEV-Pro-MTG | MKIEEGKLVIWINGDKGYNGLAEVGKKFEKDTGIKVTVEHPDKLEEKFPQVAATGDGPDIIFWAHDRFGGYAQSGLLAEITPDKAFQDKLYPFTWDAVRYNGKLIAYPIAVEALSLIYNKDLLPNPPKTWEEIPALDKELKAKGKSALMFNLQEPYFTWPLIAADGGYAFKYENGKYDIKDVGVDNAGAKAGLTFLVDLIKNKHMNADTDYSIAEAAFNKGETAMTINGPWAWSNIDTSKVNYGVTVLPTFKGQPSKPFVGVLSAGINAASPNKELAKEFLENYLLTDEGLEAVNKDKPLGAVALKSYEEELAKDPRIAATMENAQKGEIMPNIPQMSAFWYAVRTAVINAASGRQTVDEALKDAQTNSSSNNNNNNNNNNLGAMESLFKGPRDYNPISSTICHLTNESDGHTTSLYGIGFGPFIITNKHLFRRNNGTLLVQSLHGVFKVKNTTTLQQHLIDGRDMIIIRMPKDFPPFPQKLKFREPQREERICLVTTNFQTKSMSSMVSDTSCTFPSSDGIFWKHWIQTKDGQCGSPLVSTRDGFIVGIHSASNFTNTNNYFTSVPKNFMELLTNQEAQQWVSGWRLNADSVLWGGHKVFMVKPEEPFQPVKEATQLMNRRRRRGGDNGAGEETKSYAETYRLTADDVANINALNESAPAASSAGPSFRAPGSENLYFQ↓SGGDSDDRVTPPAEPLDRMPDPYRPSYGRAETVVNNYIRKWQQVYSHRDGRKQQMTEEQREWLSYGCVGVTWVNSGQYPTNRLAFASFDEDRFKNELKNGRPRSGETRAEFEGRVAKESFDEEKGFQRAREVASVMNRALENAHDESAYLDNLKKELANGNDALRNEDARSPFYSALRNTPSFKERNGGNHDPSRMKAVIYSKHFWSGQDRSSSADKRKYGDPDAFRPAPGTGLVDMSRDRNIPRSPTSPGEGFVNFDYGWFGAQTEADADKTVWTHGNHYHAPNGSLGAMHVYESKFRNWSEGYSDFDRGAYVITFIPKSWNTAPDKVKQGWPGSHHHHHH |
| MBP-TEV-Pro(K9R/  Y11A)-MTG | MKIEEGKLVIWINGDKGYNGLAEVGKKFEKDTGIKVTVEHPDKLEEKFPQVAATGDGPDIIFWAHDRFGGYAQSGLLAEITPDKAFQDKLYPFTWDAVRYNGKLIAYPIAVEALSLIYNKDLLPNPPKTWEEIPALDKELKAKGKSALMFNLQEPYFTWPLIAADGGYAFKYENGKYDIKDVGVDNAGAKAGLTFLVDLIKNKHMNADTDYSIAEAAFNKGETAMTINGPWAWSNIDTSKVNYGVTVLPTFKGQPSKPFVGVLSAGINAASPNKELAKEFLENYLLTDEGLEAVNKDKPLGAVALKSYEEELAKDPRIAATMENAQKGEIMPNIPQMSAFWYAVRTAVINAASGRQTVDEALKDAQTNSSSNNNNNNNNNNLGAMESLFKGPRDYNPISSTICHLTNESDGHTTSLYGIGFGPFIITNKHLFRRNNGTLLVQSLHGVFKVKNTTTLQQHLIDGRDMIIIRMPKDFPPFPQKLKFREPQREERICLVTTNFQTKSMSSMVSDTSCTFPSSDGIFWKHWIQTKDGQCGSPLVSTRDGFIVGIHSASNFTNTNNYFTSVPKNFMELLTNQEAQQWVSGWRLNADSVLWGGHKVFMVKPEEPFQPVKEATQLMNRRRRRGGDNGAGEETRSAAETYRLTADDVANINALNESAPAASSGGGSENLYFQ↓SGGGGSDSDDRVTPPAEPLDRMPDPYRPSYGRAETVVNNYIRKWQQVYSHRDGRKQQMTEEQREWLSYGCVGVTWVNSGQYPTNRLAFASFDEDRFKNELKNGRPRSGETRAEFEGRVAKESFDEEKGFQRAREVASVMNRALENAHDESAYLDNLKKELANGNDALRNEDARSPFYSALRNTPSFKERNGGNHDPSRMKAVIYSKHFWSGQDRSSSADKRKYGDPDAFRPAPGTGLVDMSRDRNIPRSPTSPGEGFVNFDYGWFGAQTEADADKTVWTHGNHYHAPNGSLGAMHVYESKFRNWSEGYSDFDRGAYVITFIPKSWNTAPDKVKQGWPGSHHHHHH |

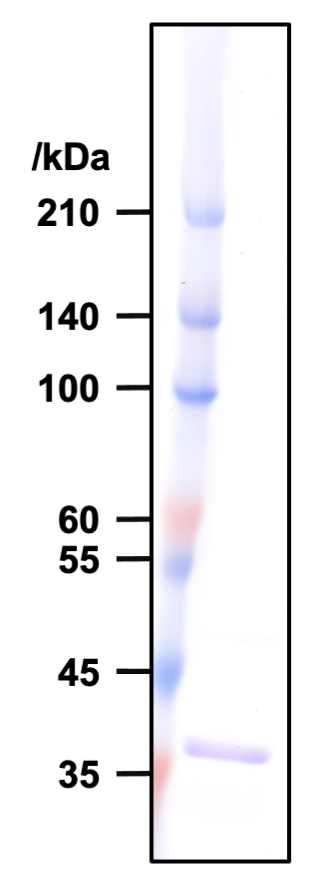

**Fig.S1. SDS-PAGE analysis of active MTG from MBP-TEV-Pro(K9R/Y11A)-MTG.** MTG sample was incubated at 37˚C for 1 h and analyzed with SDS-PAGE after heat treatment of 98˚C for 3 min in SDS-PAGE sample buffer.
